# Limelight - An open, web-based tool for visualizing, sharing, and analyzing mass spectrometry data from DDA pipelines

**DOI:** 10.1101/2024.11.01.621597

**Authors:** Michael Riffle, Alex Zelter, Daniel Jaschob, Michael R. Hoopmann, Danielle A. Faivre, Robert L. Moritz, Trisha N. Davis, Michael J. MacCoss, Nina Isoherranen

## Abstract

Liquid chromatography-tandem mass spectrometry employing data-dependent acquisition (DDA) is a mature, widely used proteomics technique routinely applied to proteome profiling, protein-protein interaction studies, biomarker discovery, and protein modification analysis. Numerous tools exist for searching DDA data and myriad file formats are output as results. While some search and post processing tools include data visualization features to aid biological interpretation, they are often limited or tied to specific software pipelines. This restricts the accessibility, sharing and interpretation of data, and hinders comparison of results between different software pipelines. We developed Limelight, an easy-to-use, open-source, freely available tool that provides data sharing, analysis and visualization and is not tied to any specific software pipeline. Limelight is a data visualization tool specifically designed to provide access to the whole “data stack”, from raw and annotated scan data to peptide-spectrum matches, quality control, peptides, proteins, and modifications. Limelight is designed from the ground up for sharing and collaboration and to support data from any DDA workflow. We provide tools to import data from many widely used open-mass and closed-mass search software workflows. Limelight helps maximize the utility of data by providing an easy-to-use interface for finding and interpreting data, all using the native scores from respective workflows.

## Introduction

Data-dependent acquisition (DDA) is the workhorse of many proteomics applications and has been the mainstream of proteomics for decades. A typical bottom-up liquid chromatography-tandem mass spectrometry (LC-MS/MS) shotgun proteomics run generates thousands of MS (MS1) spectra and tens-to-hundreds of thousands of MS/MS (MS2) spectra. Multiple tools have been developed to search DDA data to identify the peptides and pre-defined modifications that match the spectra collected.^1–4^ Recently there has been an increasing interest in “open-mass searching” of proteomics data to allow peptide-spectrum matches (PSMs) to peptides containing mass modifications that were not explicitly defined within the search parameters in advance.^5–7^ Such searches are useful when the modification is suspected but the mass of interest is unknown.^6^ This is often the case with xenobiotic-protein adducts resulting from unknown exposures, or exposure to known compounds that undergo metabolism to reactive metabolites via an uncharacterized metabolic pathway.

Many of these search tools output results using custom data formats that may work well when using that tool’s visualization software, if it exists. However, this dependency complicates data dissemination and use with external data visualization software. Some search tools do attempt to output results using standard formats, such as mzIdentML^8^ or PepXML,^9^ but even then, decisions are made by individual tool developers that affect the exact contents of these files and require knowledge of how a particular tool writes data to that standard format.

Furthermore, to generate statistics such as false discovery rates or posterior error probabilities, these tools are often used in conjunction with post-processing tools such as Percolator or PeptideProphet, that each produce still more file formats and fundamentally different kinds of scores. The sheer variety of data formats and types of scores, even from a single LC-MS/MS experiment, negatively impacts the accessibility and utility of the data. It is difficult to analyze, visualize, share, and assess the quality of the data. It is also difficult to compare output from different workflows.

Finally, different types of experiments have different visualization needs. For example, a typical closed-mass search proteomics experiment may be focused on identifying which proteins are present in given samples, while an open-mass search might be focused on discovering which modification masses are present. An investigation to determine the most appropriate set of search parameters or even which search and post-processing tools should be used, may be focused on different metrics entirely, such as how many MS features resulted in a PSM. Effective analysis and visualization of proteomics data thus requires specialized views of the data.

Several data viewers have been developed to visualize both closed-mass and open-mass search data from various analysis pipelines. Examples include closed-source commercial tools (e.g., ProteomeDiscoverer^10^), closed-source free software (e.g., MaxQuant^11^ and FragPipe^12^), and open-source free software (e.g., Trans-Proteomic Pipeline,^9^ and MetaMorpheus^7^). However, these tools are tied to specific analysis software and the output of one is often not viewable in another. The software xiSPEC^13^ is open source and does support data from virtually any DDA workflow; however, it is largely limited to viewing annotated spectra for PSMs. Additionally, most data analysis and visualization tools are not designed for secure data sharing and collaboration, nor to allow the search results and raw DDA data to be shared with the public upon publication of a specific dataset: an increasingly common requirement of proteomics journals.

We developed the software program “Limelight” to address the need for an easy-to-use, free, and open source generalized DDA results viewer, that is agnostic to the software platform used and provides an easy way to share data with collaborators or the public.^6,14^ Limelight is designed to be independent of any specific software pipeline. All its features (i.e., quality control, visualization, filtering, and analysis features) work the same, regardless of which software workflow generated the results, while simultaneously supporting visualizing and filtering using all the native scores from those software workflows. This pipeline-agnostic design is present in all aspects of Limelight, including XML schemas, database design, and web application backend and frontend code.

Limelight is a dynamic data visualization software application that is designed for 1) visualization of large proteomics data including traditional DDA (closed-mass), open-mass, and *de novo* search results and associated statistics; 2) efficient and secure data sharing either privately (collaboration within a research group) or publicly; and 3) comparing multiple experiments (e.g. control versus treated or searches of the same data with different parameters or different software workflows entirely). In addition, Limelight provides tools and metrics to visualize and analyze data quality, along with interactive statistical tools to assess data.

In this software update paper, we present the key functionality of Limelight and highlight its application using a dataset (PXD025019) comprising 6 LC-MS/MS runs from purified human serum albumin or human plasma samples which we searched with multiple software pipelines for demonstration purposes. A public Limelight project containing data used to create all visualizations presented in the following sections is available for the reader at https://limelight.yeastrc.org/limelight/p/demo.

## Methods

### Architecture

Limelight comprises multiple components, each specialized for specific tasks, linked together using a web services architecture. (Figure 1) These include a primary web application server, file object service (for storing items such as FASTA files), a web application for storing and serving spectral data, a relational database, and services for performing MS1 feature detection and spectral library generation. All components of Limelight are free and open source, developed using current software engineering best practices as follows. The front end is primarily developed using TypeScript and React (a library for developing web interfaces), standard HTML, and CSS. TypeScript is compiled into JavaScript using webpack and ts-loader as part of Limelight’s standardized build process. Limelight primarily employs a RESTful web services architecture, where the front-end client-side pages retrieve data from Limelight web services via HTTPS application programming interface (API) calls. The back-end code that handles web service calls is largely written in Java using the Spring web services framework. Limelight is built into a “.war” file suitable for hosting with Apache Tomcat (and other Servlet Containers like Jetty and WebLogic Server) using a combination of ant and Gradle.

**Figure 1.**
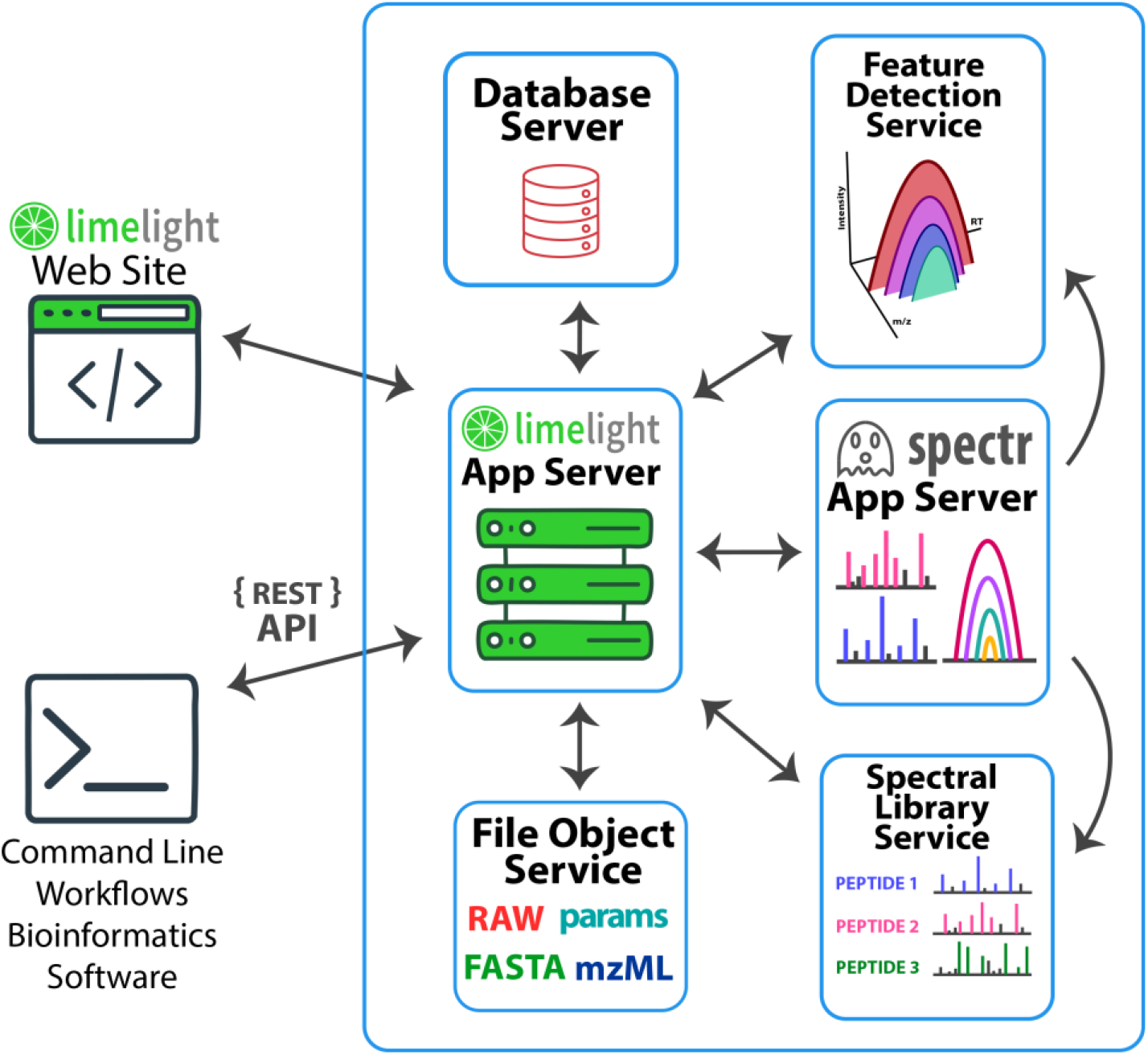
Overall software architecture of Limelight. Blue boxes around each component denote that component running in a separate Docker container. The arrows indicate which components of Limelight exchange data with each other. The surrounding blue box indicates that all components of Limelight are packaged together using Docker Compose, such that a single command will start all components and configure them correctly for intercommunication.

Limelight provides on-demand, fast access to annotated MS1 spectra, annotated MS2 spectra, and entire MS1 extracted ion chromatograms (XICs). To accomplish this, Limelight processes uploaded mzML^15^ mzXML^16^ into a binary format that is highly optimized for both minimizing storage space and retrieval of either specific scans or all scans in a range of *m/z* or retention time values. A separate web application, dubbed “spectr”, was developed to provide web-service-based API access to rapidly retrieve scan data in real time, based on user requests in Limelight. Spectr was developed in Java using a servlet framework.

Some functionality in Limelight has been implemented as external services. Examples include building spectral libraries from filtered DDA results or running MS1 feature detection. These external services are written in Python using the Flask and Flask RESTful framework and are exposed to Limelight via a RESTful web services API. The URLs for source code for Limelight, all of its components, and documentation are summarized in Table 1.

**Table 1.** URLs for Limelight source code and documentation.

| Description | URL |
| --- | --- |
| Limelight overview and launch pad | https://limelight-ms.org/ |
| Limelight user documentation and installation instructions | https://limelight-ms.readthedocs.io/ |
| Limelight server (if not installed locally) | https://use.limelight-ms.org/ |
| Limelight web application source code | https://github.com/yeastrc/limelight-core |
| Spectr source code | https://github.com/yeastrc/Spectral_Storage_Service |
| Limelight file object store service | https://github.com/yeastrc/file-object-store |
| Spectral library generation web service | https://github.com/yeastrc/limelight-export-blib-service |
| Feature detection web service | https://github.com/yeastrc/limelight-pipeline-feature-detection-service |
| Limelight XML schema definition (XSC) | https://github.com/yeastrc/limelight-import-api |

### Installation

A central Limelight server is available for users who prefer not to run it locally (Table 1). For end-user self-hosting of Limelight, each component of Limelight is distributed as Docker (https://www.docker.com/) images hosted on Docker Hub. These components are orchestrated using a Docker Compose definition file associated with a given build of Limelight on GitHub.

Docker is a framework for running pre-packaged images of software that include the software and the entire environment necessary for that software to run. Docker Compose is a component of Docker that supports the development of a relatively simple text file that describes which Docker images are used by the application and how they should be run together. To start up Limelight locally, including all of its sub-components, the end user simply needs to make changes to the configuration file and run Docker Compose. All components of Limelight will automatically be downloaded as Docker images and launched appropriately to run the entire application. When running Limelight locally, all components are run locally, and no data is sent off the individual host workstation at its premises. More information about how to run Limelight locally can be found in the user documentation and installation instructions (Table 1).

All code for all components of Limelight uses the git version control system and is hosted on GitHub (Table 1). GitHub Actions are used for continuous integration (CI) and the automated build framework. Pushes to the git repositories automatically triggers basic tests and creating a new release automatically trigger a predefined standard build procedure that builds the software, packages as Docker images, correctly tags images, pushes Docker images to Docker Hub, and associates relevant build artifacts with the respective release on GitHub.

### Data Import

Users (with sufficient privileges) can upload their search results and raw data either by using the Limelight web interface or via the command line, making it possible to integrate Limelight uploads into automated data processing workflows. The data upload section in Limelight presents users with a link for downloading a Java-based (cross platform) import submission program, documentation, and parameters necessary for authentication from the command line. When using the web interface, users select files from their computer to upload and uploads are placed into a processing queue. Data are uploaded in the form of a Limelight XML file and scan files (e.g., mzML or mzXML files). Users can see their list of pending uploads and positions in the queue as well as a log of their previous uploads (Figure 2). Once an upload has been processed, users will receive an email notification that their import was or was not successful and, if successful, the results will appear on the page as viewable in their respective sections.

**Figure 2.**
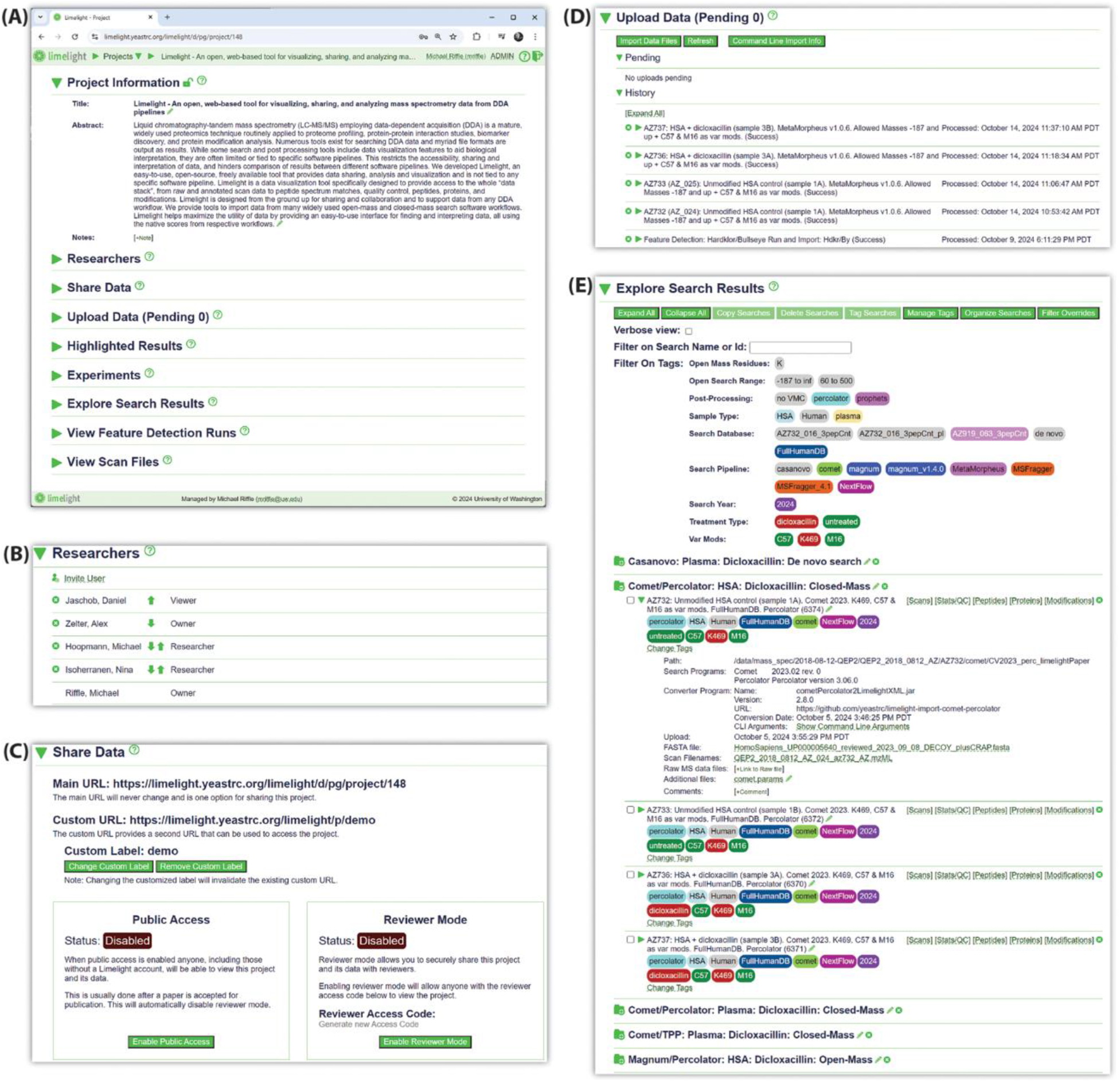
Select examples of functionality available on the project overview page in Limelight. (A) The overall structure of the project overview page. Each subsection may be expanded to access data and functionality for that section. As shown, the “Project Information” section is open, depicting the project title and abstract. (B) The “Researchers” section, which shows which users are associated with the project and provides interfaces for managing associated users and respective access level. (C) The “Share Data” section, which provides information and tools for managing the current public access level of the project. (D) The “Upload Data” section, which provides information and tools related to uploading new data to the project and reviewing logs of previously uploaded data. (E) The “Explore Search Results” section, which shows all searches uploaded to the project and their associated metadata. Tools are provided for tagging and organizing searches. Links are provided for viewing results and downloading raw data and other associated files.

Limelight Extensible Markup Language (Limelight XML) is defined by an XML Schema Definition (XSD) file (Table 1) that aims to represent the results of a DDA experiment in a generalized format that is pipeline-agnostic and contains all data necessary for Limelight. It encodes many general aspects of a DDA search and any post processing tools, including which search programs were used, encoded configuration files, scores associated with PSMs, peptide-level scores, protein-level scores, modification-level scores and peptide-protein mappings. Additionally, Limelight XML encodes which scores are present in the file for each of the software programs used, what those scores are describing (e.g., PSMs or peptides), and characteristics of those scores (e.g., smaller numbers are better). Limelight does not make any assumptions about the data beyond what is represented in the Limelight XML document.

DDA results from any workflow that are represented as Limelight XML may be imported into Limelight and visualized using all tools within Limelight. This moves the responsibility of decoding and staying current with the output of specific workflows to individual software programs, here called Limelight XML converters, that will convert native output from software pipelines to Limelight XML. These Limelight XML converters can be created and maintained independently of Limelight itself, which reinforces the pipeline-agnostic design. The authors have written and maintain Limelight XML converters for many popular DDA software workflows (Table 2) and will continue to expand the list of available converters by directly developing them or in collaboration with authors of LC-MS/MS search software.

**Table 2.** Current list of Limelight XML converters built and maintained by the authors.

| Software Pipeline | Limelight XML converter, source code, and documentation |
| --- | --- |
| Casanovo (*de novo* AI)<sup>17</sup> | https://github.com/yeastrc/limelight-import-casanovo |
| Comet<sup>2</sup> + Percolator<sup>18</sup> | https://github.com/yeastrc/limelight-import-comet-percolator |
| Comet + Trans Proteomic Pipeline<sup>19</sup> | https://github.com/yeastrc/limelight-import-comet-tpp |
| Comet-PTM<sup>20</sup> | https://github.com/yeastrc/limelight-import-cometptm |
| Crux<sup>21</sup> | https://github.com/yeastrc/limelight-import-crux-comet-percolator |
| Magnum<sup>6</sup> + Percolator | https://github.com/yeastrc/limelight-import-magnum-percolator |
| Magnum + Trans Proteomic Pipeline (TPP) | https://github.com/yeastrc/limelight-import-magnum-tpp |
| MetaMorpheus<sup>7</sup> | https://github.com/yeastrc/limelight-import-metamorpheus |
| modA<sup>22</sup> | https://github.com/yeastrc/limelight-import-moda |
| MSFragger<sup>5</sup> | https://github.com/yeastrc/limelight-import-msfragger-tsv |
| MSFragger + TPP | https://github.com/yeastrc/limelight-import-msfragger-tpp |
| Open pFind<sup>23</sup> | https://github.com/yeastrc/limelight-import-open-pfind |
| Philosopher<sup>24</sup> (MSFragger or Comet) | https://github.com/yeastrc/limelight-import-philosopher-tsv |
| ProLuCID<sup>25</sup> (DTASelect,<sup>26</sup> using IP2 | https://github.com/yeastrc/limelight-import-prolucid-dtaselect |
| Tag Graph<sup>27</sup> | https://github.com/yeastrc/limelight-import-taggraph |

### Database Design

Limelight is built using the MySQL relational database management system (RDBMS). Like the Limelight XML schema described above, the database schema is designed to store DDA search metadata and results in an agnostic, generalized way. No *a priori* knowledge of any software workflow or types of scores are built into the database design. Tables and relationships exist to store the annotations (e.g., scores) associated with various types of data in DDA results (e.g., PSMs) in a way that is both generalized and optimized for searching, so that the native scores in the respective workflows can be viewed, efficiently searched, and used for filtering results in Limelight.

## Results and Discussion

### Project Interface

All data in Limelight are organized into projects (Figure 2). From the project page, users may (depending on their access level) add or change metadata, control who has access to the project (Figure 2B), upload data (Figure 2D), run simple workflows, download raw data, or navigate to a page to view search results (Figure 2E). Projects can be limited in scope (containing a handful of searches) or extensive (containing dozens or hundreds of searches). For projects containing many searches, organizing searches becomes critical to efficiently and productively using Limelight.

Limelight provides two primary mechanisms for organizing search results: folders and tags. Users can create folders with meaningful names within a project and organize their searches under those folders. Tags and tag categories are defined by users at the project level, allowing users to create a simple customized controlled vocabulary for labeling searches in the project (Figure 2E). For example, a tag category called “Sample type” could be created with tags for “Human”, “HSA”, and “plasma” using different colors. Searches can then be labeled with tags belonging to each of these categories, marking them visually with a simple ontology that is custom for the project. Tags are shown when searches are listed, giving an effective means of rapidly assessing which searches are relevant to the current user. Additionally, these tags may be used as filters on the project page to quickly find relevant searches.

The “Explore Search Results” section (Figure 2E) of the project page lists all search results uploaded to the project. Searches are listed with user defined names and tags and are organized by folder (if any are defined). Users may view metadata for each search or download and view raw, FASTA, configuration, or other files associated with the search. Each search listing includes links for viewing the results using specialized viewers that emphasize a different level of data: scan-by-scan, peptide-level, protein-level, and modifications-level (e.g., PTMs).

Additionally, a Stats/QC viewer provides many visualizations summarizing various aspects of the search results.

### Data Visualization Tools

As previously highlighted,^6^ the peptide, protein and modification level visualization views in Limelight allow the user to explore the data from a peptide-, protein-, or modification-centric point of view (Figure 3). Each of these views allows drilling down to the peptide and PSM level data with all the native, pipeline-specific scores displayed. Each PSM also includes a link to view the annotated MS2 spectrum using Lorikeet^28^ (Figure 3B) and to view the annotated MS1 spectrum for the precursor, highlighting the expected isotopic peaks. Additionally, users may view the XIC for the peptide (Figure 3A) in the context of a given search, which shows where PSMs were identified for the given peptide. The XIC viewer interacts with spectr (described above) to construct the XIC in real time.

**Figure 3.**
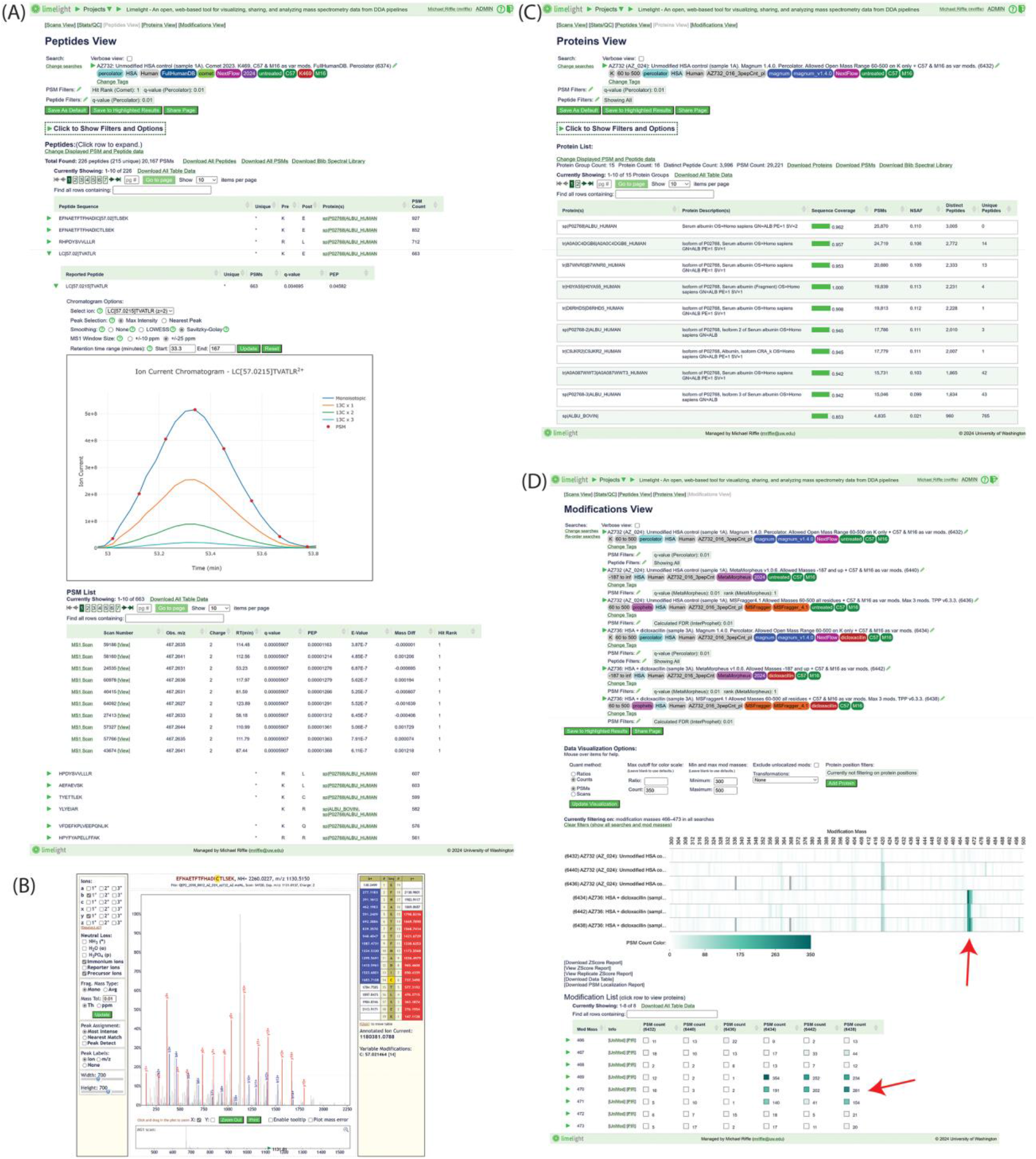
Select data views from Limelight. (A) The “Peptides View” is a peptide-centric view of the data. All peptides (that pass current filter options) are listed, and each may be expanded for more information. In this example, “LC[57.0215]TVATLR” is expanded. All native peptide- and PSM-level scores are available. An XIC for the peptide is shown with red points indicating at what retention time and isotopic peak a PSM was sampled for this peptide. (B) Annotated MS2 spectrum as shown by Limelight. All b+ fragment ions are shown in blue and y+ fragment ions are shown in red. (C) The “Proteins View” is a protein-centric view of the data. All protein groups are shown as rows, including any scores and other data (e.g., sequence coverage, number of PSMs, and so forth) associated with that protein group. Each protein may be clicked for more information. (D) The “Modifications View” is a PTM (or other modification) centric view of the data, focusing on which modification masses were observed in the data. In this example, one sample each for untreated and dicloxacillin-treated purified human serum albumin (HSA) was searched using three separate open-mass software pipelines (MetaMorpheus, MSFragger + TPP, and Magnum + Percolator) and compared using the modification view in Limelight. The heatmap indicates the frequency of observations of PSMs with a given modification mass (rounded to the nearest integer) in each search for all modification masses observed (filtered to 300-500 Da for this example). The three top rows are controls, and the three bottom rows are the treated samples. The red arrow indicates the expected modification mass for the dicloxacillin treatment. The table below provides an interface for browsing all observed modification masses and their associated proteins, peptides, and PSMs. The red arrow indicates the rows associated with the expected modification mass of dicloxacillin treatment. (See: https://limelight.yeastrc.org/limelight/go/OY5VgsQaHs)

The protein view (Figure 3C) allows users to control protein inference (protein grouping).

By default, proteins are grouped using a protein parsimony algorithm that uses a greedy set cover algorithm to estimate the fewest number of protein groups necessary to explain the observed peptides. Other options include no grouping, only including groups that are not subsets of other groups, or, including all groups. The protein groups are displayed in rows in the viewer and the rows may contain more than one protein if grouping is enabled. Additionally, users may choose to show computationally derived columns for each protein that include normalized spectrum abundance factor (NSAF)^29^ adjusted spectral counts using the algorithm described by ABACUS^30^ and the NSAF calculated using adjusted spectral counts.

To provide more data integrity visualization, a new primary data visualization page called the “Quality Control (QC) view” has been added to Limelight. This page includes numerous tables and data visualizations describing various characteristics of the data that are separated by category. For a single search (Figure 4), categories include summary statistics, digestion statistics, target-decoy statistics, feature detection statistics, scan file statistics, chromatography statistics, PSM-level statistics (error estimates, distributions of PSM counts and scores, and relationships between scores), and peptide level statistics.

**Figure 4.**
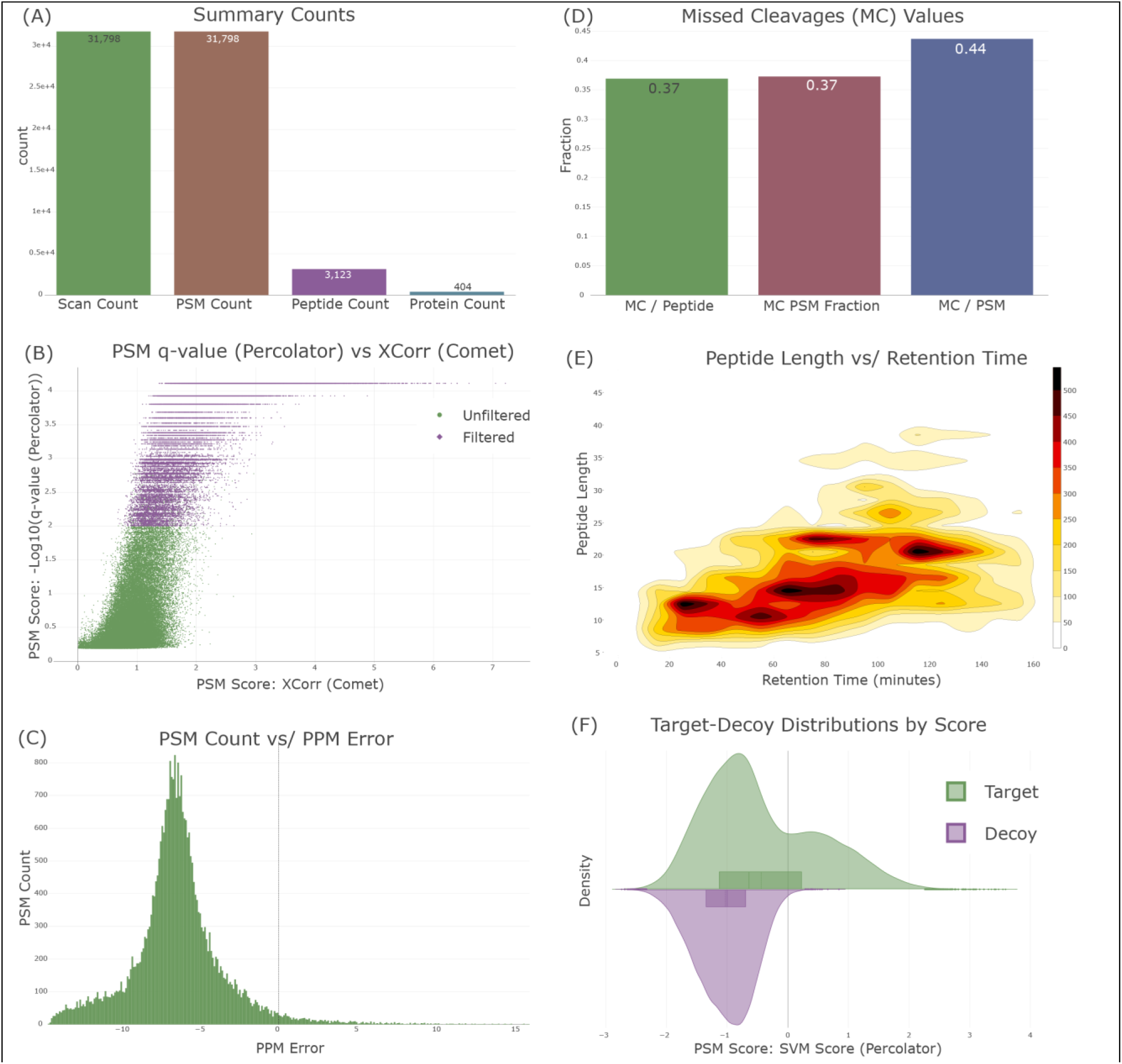
Select examples of visualizations available on the “QC View” in Limelight. (A) The “Summary Counts” visualization indicates the number of distinct scans, PSMs, peptides, and proteins that were observed in the results. (B) An example of plotting PSM-level scores in the results against any other PSM-level score. In this case, the -log10 of the Percolator q-value is plotted against the Comet Xcorr (cross correlation) score for the PSMs found in the given search. All PSMs passing the current filters (labeled “Filtered”) shown in purple and the rest shown in green. (C) A histogram of the ppm error associated with observed PSMs. (D) A plot indicating the number of missed cleavages per peptide, the fraction of PSMs containing a missed cleavage, and the number of missed cleavages per PSM. (E) A 2D density plot illustrating the relationship between peptide length and retention time, based on observed PSMs. (F) A plot that shows the distributions of targets and decoys for any PSM-level score in the results. In this case the Percolator SVM score is shown.

When comparing multiple searches in the QC view, many of the plots will change from the single-search view to one that emphasizes the differences between multiple searches using the displayed metrics (Figure 5). For example, the summary statistics and digestion statistics plots change to show each statistic in a separate plot that compares those statistics across searches. Nearly all plots are available for comparing searches, though some (e.g., chromatography and target/decoy statistics) are not appropriate for comparing searches and are not displayed.

**Figure 5.**
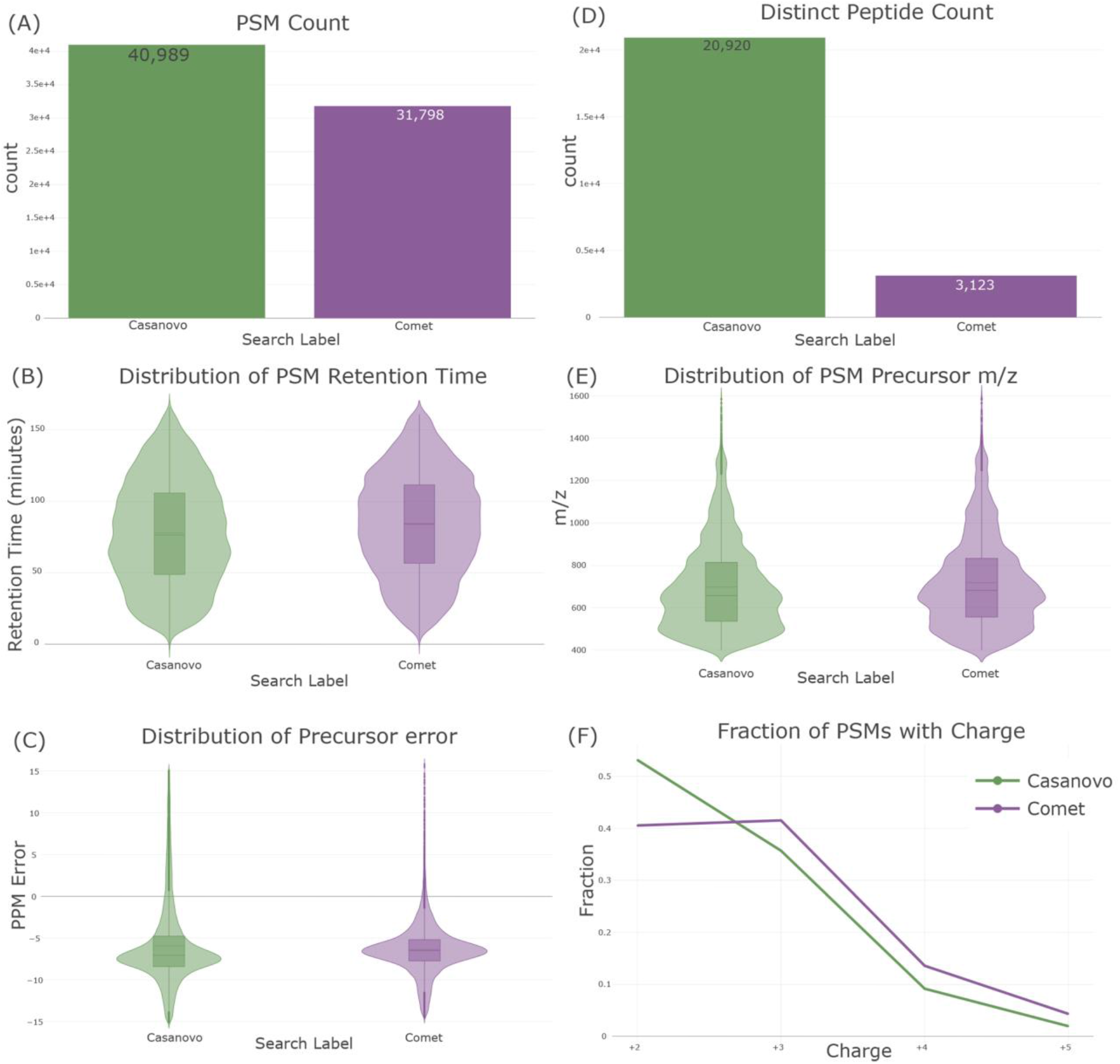
Select examples of comparing multiple searches using the “QC View” in Limelight—in this case, comparing a Casanovo *de novo* search (green) with a comet search (purple) of the same raw data. (Note that this is not meant to be a comparison of these two workflows, but only illustrative of views in Limelight.) (A) A comparison of the number of PSMs observed in the two workflows. (B) A comparison of the distribution of retention times. (C) A comparison of the distribution of precursor PPM errors. (D) A comparison of the number of observed distinct peptides. (E) A comparison of the distribution of precursor *m/z* values. (F) A comparison of the number of observed PSMs with a given charge in the two workflows.

Many of the plots on the QC view are interactive and dynamic. Clicking on the plot will open that plot in a larger window and options will appear for customizing (e.g., changing which statistics are plotted or zooming in) and downloading that plot as a vector graphic image (SVG) or bitmap image (PNG).

### Filtering Data

Data filtering is one of Limelight’s most powerful features and is a prominent part of every data view page described. The filtering options allow users to drill down to biologically interesting proteins, peptides, modifications or other data features of interest in large and complex datasets. Every page that displays search results will display the list of searches being viewed at the top of the page (Figure 3), as well as any filters that are being applied using the scores in those searches. Any native scores associated with the specific pipeline can be used to filter PSM, peptide, protein, or modification results. Many search workflows have a default filter value for their results (as defined in the Limelight XML file that was imported) automatically applied – for example, a filter of 0.01 might be applied to all Percolator PSM-and peptide-level q-values if that was specified as a default cutoff in the Limelight XML. The filters being applied may be changed on the web interface and the page automatically updates using the new filter values. Limelight solely uses the description of the scores from the Limelight XML to determine how to filter on specific scores (e.g., smaller is better).

In addition to search-based score filters, more generalized filter options exist for pages where they are applicable. For example, peptides can be filtered on precursor *m/z* or retention time ranges, precursor charge(s), PSM counts, specific modification masses, and so on.

Proteins can be filtered on minimum PSM counts, minimum peptide counts, and a minimum number of peptides that map specifically to that protein. In the single protein view, the sequence coverage interface can also be used as a filter by clicking one or more residues to filter for peptides covering those positions.

### Comparing and Contrasting Search Results

An important feature of Limelight is that it allows side-by-side comparison and visualization of multiple experiments (Figure 3D and Figure 5). This feature has been used extensively to compare peptide modifications resulting from drug treatment to untreated control samples in both open-mass search^6^ and closed-mass search^14^ workflows. Limelight also supports comparing results from different DDA search pipelines in the same interface and makes no assumptions about which workflows generated each dataset. This means that Limelight supports comparing results from different software workflows (or different versions of the same workflow) in a single web interface, using the native scores from each set of results. All the features of the web application will function in a consistent way, regardless of the workflow that generated the data–so long as it can be represented by Limelight XML. All the data view pages described above can be used for single searches or with multiple searches. When viewed with multiple searches, the views may change to accommodate displaying scores and statistics from multiple searches, instead of one. Viewing multiple searches is simple; the user selects the searches they would like to compare and clicks the relevant “Compare” button in the “Explore Search Results” section of the project page.

### Feature Detection

MS1 feature detection is the identification of persistent, peptide-like isotope distributions observed in consecutive MS1 scans that indicate where in time peptides may be eluting–even if they are not observed as PSMs. Limelight provides tools for users to run a Hardklör^31^ and Bullseye^32,33^ feature detection workflow, view the predicted features, and view those features in the context of search results that have been uploaded for the same scan file (Figure 6).

**Figure 6.**
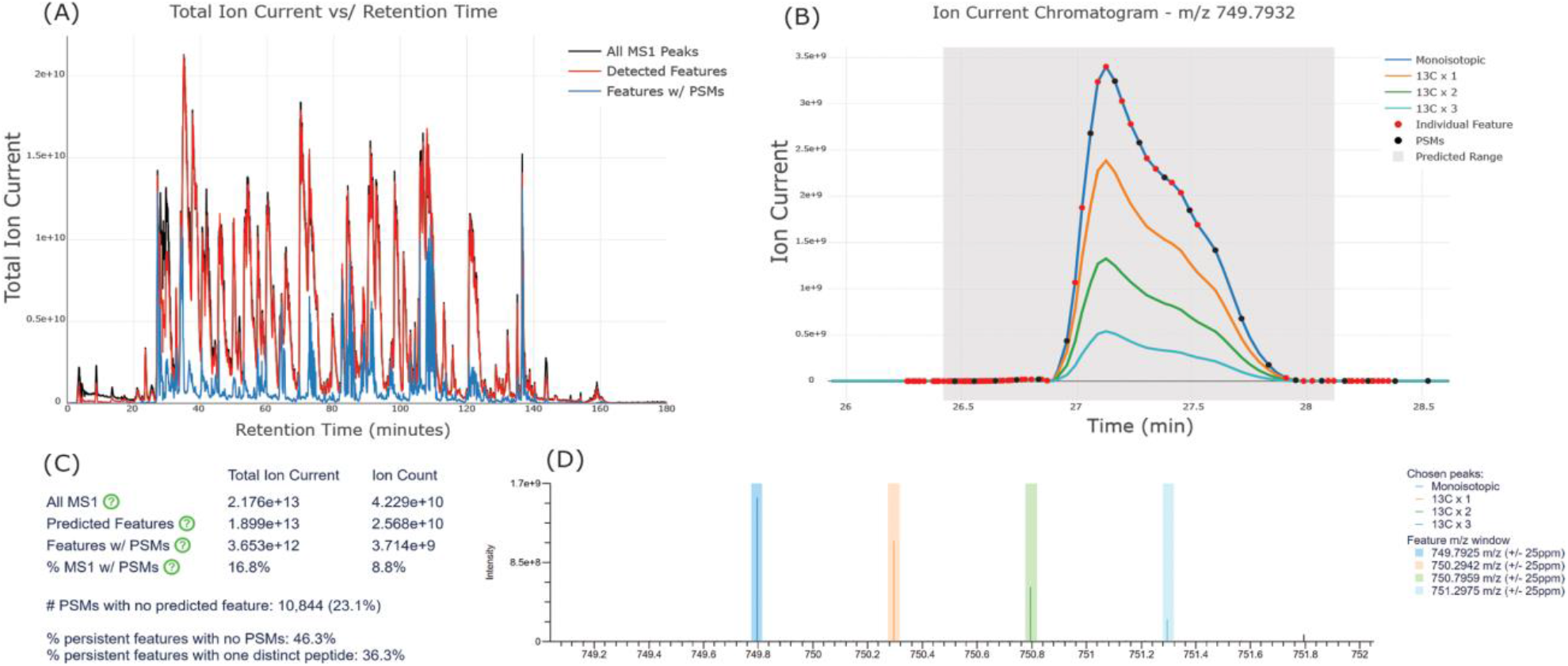
Select examples of data and visualizations resulting from running MS feature detection in Limelight. (A) A plot of the total ion current versus retention time for all MS1 peaks (black), all detected MS1 features (red), and all PSMs that correspond to detected features (blue). (B) XICs are available for predicted persistent features. Chromatograms for 4 isotopic masses are shown. The gray shaded region indicates the predicted retention time range, the red balls indicate individual MS1 features used to predict the persistent feature, and the black balls indicate PSMs in the data for the given retention time range and given *m/z* values. The red and black balls may be moused over or clicked for more information, such as what the peptide identifications were for the PSMs. (C) A table indicating summary information, such as total MS1 ion current (TIC), the TIC associated with predicted features, the TIC associated with features associated with PSMs, and the fraction of the MS1 TIC associated with a feature that is associated with a PSM. Values are also shown for ion count, calculated using the ion current (ions per second) with the ion injection time for each scan. Additionally, the number of PSMs with no predicted feature, the number of features with no PSM, and the number of persistent features that map to exactly one distinct peptide are shown. (D) The annotated MS1 scans for any predicted feature may be viewed. The colored areas indicate the expected location of the isotopic peaks.

To run a feature detection workflow, project owners expand the “View Scan Files” section on the project page, which displays all scan files that have been uploaded to the current project (usually done when search results are uploaded). The user then clicks the “Run Feature Detection” link for a given scan file, supplies configuration details, and runs the workflow. Once done, the user is notified of completion and may view the feature detection results from the project page or from the QC page for search results that used that scan file.

Feature detection results can also be viewed from the project page by either expanding the “View Feature Detection Runs” section or by expanding the “View Scan Files” section and then the scan file of interest. When viewing a feature detection run, users are presented with data visualizations related to how much of the total ion current (TIC) is covered by the predicted features, a histogram showing the distribution of features by TIC, and a list of all persistent features predicted by Bullseye (Figure 6). Data from Bullseye, such as charge, monoisotopic mass, retention time range, and abundance are shown in tabulated format for all features. Each identified feature may be expanded to view the associated XIC and list of Hardklör predictions used to assign the persistent feature. Additionally, search results may be loaded for the given scan file that adds statistics and visualizations related to how many of the features have observed PSMs, how many PSMs have predicted features, how many features map to a single predicted peptide, and how TIC from PSMs compares to TIC of predicted features.

### Data Sharing, Security and Collaboration

Limelight is designed to be a secure data-sharing platform–both for internal collaboration and for public sharing. All access to data is secured by controlling access to projects, and by default, all data uploaded to a project are private, and only users associated with that project have access to data in that project. This “private” level of sharing, where the owner of the project can optionally share access with one or more specific Limelight user, is intended to be used for private data sharing and collaboration. To share data with other users of Limelight, project owners may add existing Limelight users or invite users to create a Limelight account and join the project. When inviting non-Limelight users, Limelight sends an email that contains a special link for creating an account and joining the project. Project owners may also remove users from the project (removing their access to data), or change access levels (e.g., moving a user from read-only to read/write).

To share data with the public (viewers will not need a Limelight account) two options exist. First a project can be marked as “public”—anyone accessing the project’s unique URL will have access to the project and its data (Figure 2C). For example, the project associated with this publication (https://limelight.yeastrc.org/limelight/p/demo) is a public project. This public sharing mode is intended to be used after a manuscript is accepted for publication or if data are generated that the user wishes to make public. Public mode can be enabled or disabled at any time by the project owner. Second, “reviewer mode” may be enabled for a project (Figure 2C). Reviewer mode generates a short, unguessable random string that a non-Limelight user can supply when accessing a project that grants them read-only access to the project and its data (along with appropriate items like the project title and abstract). This is intended to be used when submitting a manuscript so that reviewers may access the data without the need to make the data public. Reviewer mode can be disabled at any time and the random string can be regenerated, which invalidates the existing string.

Project owners may also customize the URL associated with a project to be more descriptive. Additionally, the project owner can lock a project, which prevents any further changes to items like the title, abstract, or data associated with the project.

### Usage and Scalability

Limelight is designed to be performant and responds well to both vertical and horizontal scaling. The main web application, database, spectr (storing and serving spectra data), and web services for feature detection, spectral library generation, and file object storage (storing files such as FASTA files) are encapsulated and can be run either on the same system or separate systems. Special attention has been paid to efficiency of the code in the web application, schema of the relational database, storage architecture, and binary formats used by spectr to ensure performance. Increasing the capabilities of the CPUs, amount of RAM, and performance of the storage media can have large impacts on the performance of Limelight and its components. Likewise, moving services (such as the web application, database, and spectr) to separate servers can improve performance.

The authors host an internal installation of Limelight that is used regularly by researchers for a variety of analysis tasks and has remained performant. This installation hosts the web application and database on a single server with an Intel Xeon CPU (from 2016), 256 GB RAM, and a NVMe solid state disk. As of this writing, this system hosts 106 users, 5,287 MS/MS searches, nearly 68 million peptides identified in those searches, and nearly 487 million PSMs. Spectr is hosted on a separate server that hosts data from 8,284 scan files taking up 4.7 TB of space using its binary format and indices.

Documentation and channels for requesting support are critical features of a web application as full featured as Limelight. Limelight contains a large amount of inline help via tooltips that appear when mousing over special help icons. Additionally, online documentation exists at https://limelight-ms.readthedocs.io/ describing installation, administration, tutorials, and basic usage of Limelight. For support, users and developers may submit issues via GitHub (https://github.com/yeastrc/limelight-core/issues), join our Slack at https://limelight-ms.slack.com/, or send an email to the address listed at the bottom of the site.

### Limitations and Future Directions

The chief limitation of Limelight is a lack of native support for label free quantification (LFQ) pipelines, MS1 peak area, and isobaric labeling workflows using tandem mass tags (TMTs). Additionally, Limelight is currently limited to DDA workflows. Technologies like data-independent acquisition (DIA), parallel reaction monitoring (PRM), and selected reaction monitoring (SRM) are well established mass spectrometry-based methods for protein quantification. Future work in Limelight will be to integrate DDA-based quantification methods into the existing schema and user interface models, build converters for such workflows, and to build a generalized model to support protein quantification workflows that encapsulate DIA, PRM, and SRM peptide-centric workflows (i.e., a peptide-centric Limelight XML schema) to present these results to end users in a biological context.

## Acknowledgments

The work presented here was supported by the National Institutes of Health, National Institute of General Medical Sciences under Award R01GM147947 (to N.I.) and P41GM103533 (to M.J.M.), and the National Institute on Aging under Award Numbers R01GM087221, S10OD026936, and from the National Science Foundation Grant DBI-1933311 (to R.L.M.). Support was also provided by the Intelligence Advanced Research Projects Activity (IARPA) TEI-REX program through the Army Research Office contract W911NF2220059. The views and conclusions contained should not be interpreted as necessarily representing the official policies, either expressed or implied, of ODNI, IARPA, ARO, or the U.S. Government. The U.S. Government is authorized to reproduce and distribute preprints for governmental purposes notwithstanding any copyright annotation therein. This work was supported in part by the University of Washington’s Proteomics Resource (UWPR95794).

## Notes

https://limelight-ms.org/

### Competing Interest Statement

The MacCoss Lab at the University of Washington receives funding from Agilent, Sciex, Shimadzu, Thermo Fisher Scientific, and Waters to support the development of Skyline, a quantitative analysis software tool. MJM is a paid consultant for Thermo Fisher Scientific.

